# *In silico* characterization and expression profiles of zinc transporter-like (LOC100037509) gene of tomato

**DOI:** 10.1101/2020.10.03.324913

**Authors:** Ahmad Humayan Kabir

## Abstract

Zinc (Zn) is an essential microelement for plants. ZIP transporters play a critical role in Zn homeostasis in plants. This *in silico* study characterizes different features of putative Zn transporter of tomato (Solyc07g065380) and its homologs. A total of 10 ZIP protein homologs were identified across nine plant species by protein BLAST. All these ZIP protein homologs located at chromosome 7 showed 305-350 amino acid residues, 7-8 transmembrane helices, and stable instability index. Further, these ZIP protein homologs are localized in the plasma membrane at the subcellular level corresponding to the ZIP zinc transporter (PF02535) domain. Gene organization analysis reveals the presence of 3 exon along with the position of the promoter, TATA-box, transcriptional start site, and splice sites in these ZIP transporter homologs, in which tomato ZIP transporter (NM_001247420.1) contains a promoter, TATA-box, transcriptional start site at 500, 911 and 946 bp, respectively along with several splice sites, which may be useful for targeting binding sites and transcription factor analysis. Further, the cutting sites and restriction enzymes of each ZIP gene homologs might be helpful for future transgenic studies underlying Zn homeostasis. MEMO displayed five conserved motifs associated with the ZIP zinc transporter, N-glycosylation site, and phosphorylation site. Phylogenetic studies reveal a close relationship of Solyc07g065380 with *Solanum pennellii* homolog, while ZIP transporter of *Nicotiana sylvestris* and *Nicotiana tabacum* predicted to be in close connection. The Solyc07g065380 transporter is predominantly linked to several uncharacterized zinc metal ion transporters and expressed in diverse anatomical part, developmental stage, and subjected to pathogen and heat stress. The secondary structural prediction reveals unique signal peptide in the ZIP protein homologs of *S. lycopersicum* and *S. pennellii* along with extended alpha-helix. These bioinformatics analyses might provide essential background to perform wet-lab experiments and to understand Zn homeostasis for the development of Zn-biofortified crops.

**Key message:**

♦ ZIP protein homologs are localized in the plasma membrane and are linked to ZIP zinc transporter (PF02535) domain at chromosome 7.
♦ ZIP protein motifs are associated with the ZIP zinc transporter, N-glycosylation site, and phosphorylation site.
♦ Phylogenetic studies reveal a closet relationship of Solyc07g065380 with *Solanum pennellii* homolog.
♦ ZIP protein homologs of *S. lycopersicum* and *S. pennellii* show unique signal peptide along with extended alpha-helix.

## Introduction

Zinc (Zn) shortage, a plant nutritional deficit, has adverse effects on vegetable (Mattiello et al. 2015). There are lands with high pH or alkaline stress in many parts of the world (Alloway 2009; Cakmak 2008), the primary explanation for Zn-deficient soil. The soil of lower pH, sandy texture, and lower phosphorus are often deficient in Zn (Marschner, 1995). Zn-starved plants often show impressive growth, tiny stems, and mild chlorosis (Marschner, 1995). Zn functions in photosynthetic and gene expression processes in in addition to enzymatic and catalytic activities (Welch 2001). Zn deficiency also resulted in a decline in plant stomatal activity (Mattiello et al. 2015). In response to pathogenic attacks on plants, Zn proteins also play a significant role (Cabot et al. 2019). Zn’s insufficiency in plant-based foods contributes to global human malnutrition (Hotz et al. 2004).

While generally there are genotypic disparities in Zn-deficiency responses, lots of plants are susceptible to Zn deprivation (Hacisalihoglu et al. 2001; Khatun et al. 2018). While Zn efficiency mechanisms are fairly complex, research found several adaptive characteristics in Zn deficient plants (Höller et al. 2014; Hacisalihoglu et al. 2003). As Zn cannot penetrate the root cell membranes, Zn’s absorption is achieved primarily via transporters (Kabir et al. 2017). The *IRT*, known as Fe-regulated transporter is often induced to enhance translocation as well as Zn absorption whenever the plant has possibly Zn or Fe shortcomings (Lin et al. 2009; Eckhardt et al. 2001). *IRT1* is mainly responsible for transporting different metals like Zn, Mn, and Fe, despite the fact that the pattern of expression varies depending on mineral stresses (Durmaz et al. 2011; Eckhardt et al. 2001). Till today, progress is pronounced to imparting the performance of *ZIP* (Zn transporter) in Fe and Zn uptake (Morea et al. 2002; Xu et al. 2010), though the proof in tomato continues to be restricted. Pavithra et al. (2016) classified low-affinity (*SlZIP5L2, SlZIP5L1, SlZIP4*, and *SlZIP2*) and high-affinity (*SlZIPL*) Zn transporters in tomato. In a recent study (Akther et al. 2020) demonstrated that upregulation of SlZIPL and SlZIP2 is involved with the increase of Zn in Zn-deficient tomato. However, there is a putative zinc transporter (Solyc07g065380) provisionally submitted to NCBI.

Tomato (*Solanum lycopersicum* L.) is a high-value crop worldwide (Wilcox et al., 2003). The deficiency of the essential micronutrient Zn harms tomato production worldwide. Therefore, the characterization of Zn transporter possesses immense potential to combat Zn-deficiency in plants. Therefore, the molecular characterization of ZIP homologs may provide in-depth insight into these genes/proteins. In this study, we have searched for ZIP transporter homologs based on putative zinc transporter of tomato (Solyc07g065380/LOC100037509) in different plant species. The CDS, mRNA, and protein sequences of these ZIP homologs were taken into different computational analysis with advanced bioinformatics and an online-based platform along with the wet-lab expression of mRNA transcript of putative zinc transporter in tomato under differential Zn-conditions.

## Materials and Methods

### Retrieval of ZIP genes/proteins

Zn transporter-like (LOC100037509) gene named as Solyc07g065380 in Uniprort/Aramene database (protein accession: NP_001234349.1 and gene accession: NM_001247420.1) was obtained from NCBI to use as a reference for homology search (Stephen et al. 1997). The search is filtered to match records with expect value between 0 and 0. The corresponding FASTA sequences of gene and protein were retrieved from NCBI database.

### Analyses of ZIP genes/proteins

Physico-chemical features of MTP protein sequences were analyzed by the ProtParam tool (https://web.expasy.org/protparam) as previously described. Transmembrane (TM) helix prediction was carried out by TMHMM Server (http://www.cbs.dtu.dk/services/TMHM). The CELLO (http://cello.life.nctu.edu.tw) server predicted the subcellular localization of proteins (Yu et al. 2006). Chromosomal locations, protein domain families, and functions were searched in ARAMEMNON (http://aramemnon.uni-koeln.de/).

### Organization of ZIP genes

The organization of exon and intron was analyzed by the ARAMEMNON web tool. Organization of coding sequence along with the position of the transcriptional start site, TATA-box, and the splice sites were predicted by FGENESH, TSSPlant, and FSPLICE tools (http://www.softberry.com/berry.phtml). Further, the promoter site was predicted by the Promoter 2.0 Prediction Server (http://www.cbs.dtu.dk/services/Promoter/). The analysis of restriction sites and enzymes were predicted by Restriction Analyzer (http://molbiotools.com/restrictionanalyzer.html) and Restriction Enzyme Picker Online v.1.3 (https://rocaplab.ocean.washington.edu/tools/repk).

### Phylogenetic relationships and identification of conserved protein motifs

Multiple sequence alignments of MTP1 proteins were performed to identify conserved residues by using Clustal Omega and MView. Furthermore, the five conserved protein motifs of the proteins were characterized by MEME Suite 5.1.1 (http://meme-suite.org/tools/meme) with default parameters, but five maximum numbers of motifs to find. Motifs were further scanned by MyHits (https://myhits.sib.swiss/cgi-bin/motif_scan) web tool to identify the matches with different domains (Sigrist et al. 2010). The MEGA (V. 6.0) developed the phylogenetic tree with the maximum likelihood (ML) method for 1000 bootstraps using ZIP homologs (Tamura et al. 2013).

### Interactions and co-expression of tomato ZIP protein

The interactome network of Solyc07g065380 was generated using the STRING server (http://string-db.org) visualized in Cytoscape (Szklarczyk et al. 2019). Further, gene co-occurrence and neighborhood pattern were also retried from the STRING server. Additionally, the expression data of Solyc07g065380 was retrieved from Genevestigator software and analyzed at hierarchical clustering and co-expression levels on the basis of the SL_mRNASeq_Tomato_GL-0.

### Structural analysis of ZIP protein homologs

Analysis of the outer membrane of ZIP proteins was performed by Philius Transmembrane Prediction Server (http://www.yeastrc.org/philius/pages/philius/runPhilius.jsp) as previously described (Reynolds et al. 2008). The secondary structure was described by the SOPMA tool (https://npsa-prabi.ibcp.fr/cgi-bin/npsa_automat.pl?page=npsa_sopma.html).

## Results

### Retrieval of Zn transporter-like genes/proteins

The blast analysis of the Zn transporter-like (LOC100037509) gene of tomato showed 10 homologs by filtering the E-value to 0.0. The retrieved gene/proteins include *Solanum lycopersicum, Solanum pennellii, Solanum tuberosum, Nicotiana sylvestris, Nicotiana tabacum, Capsicum annuum, Nicotiana tomentosiformis, Nicotiana attenuate* and *Camellia sinensis* (Table 1).

**Table 1.**
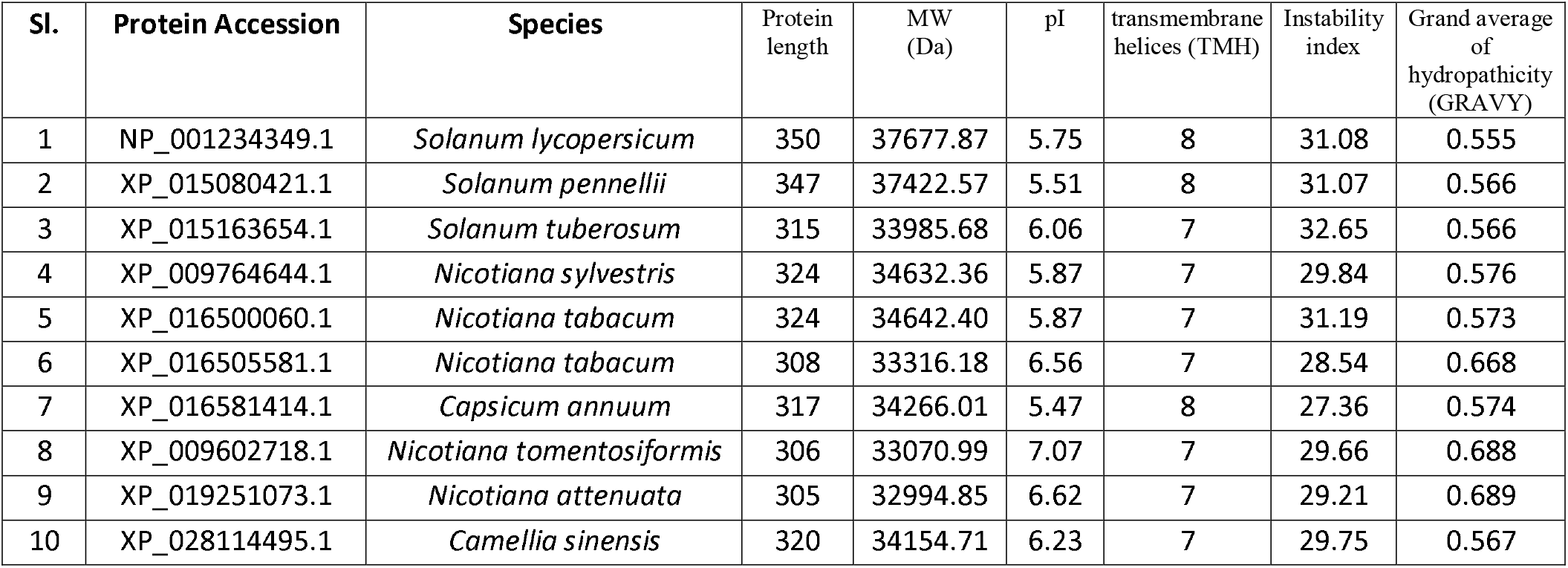
Physiochemical properties of ZIP transporter homologs.

### Physiochemical features and localization of MTP proteins

In total, 10 ZIP homolog proteins encoded a protein with residues of 305–350 amino acid having residues having 32994.85-37677.87 (Da) molecular weight. The pI value, instability index, and Grand average of hydropathicity ranged from 5.51-7.07, 27.36-31.19 (stable), 0.555-0.689, respectively (Table 1). Notably, all these ZIP proteins showed 7-8 transmembrane helices (TMH). All of these ZIP protein homologs belonging to the ZIP zinc transporter (PF02535) domain localized in the plasma membrane having membrance as cellular components (Table 2). These ZIP proteins are located in chromosome 7 involved in metal ion transmembrane transporter activity (Table 2).

**Table 2.**
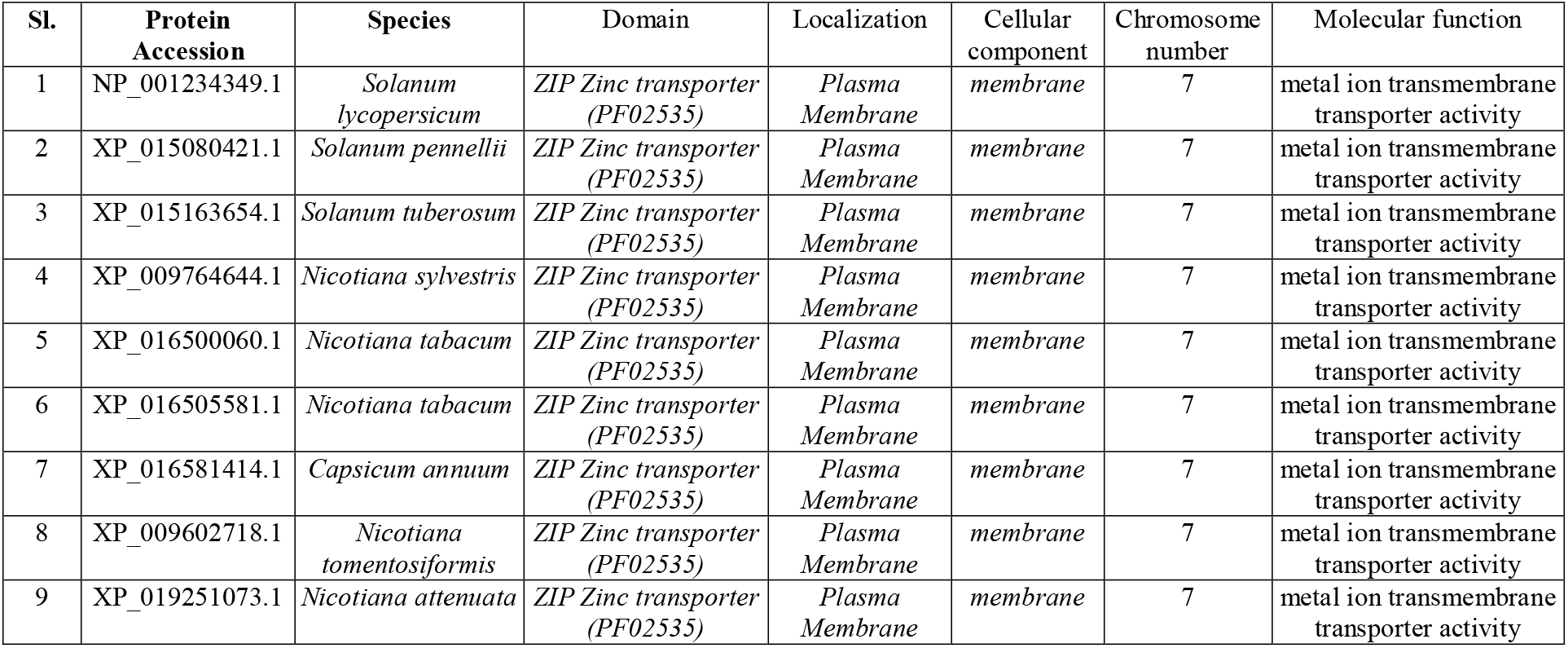
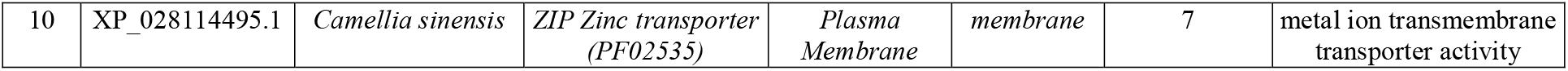
Domain, localization, cellular component, chromosome position and molecular function of ZIP proteins.

### Organizational features of ZIP transporter genes

The structural analysis of the ZIP genes showed the presence of 1 exon the coding sequence, which varies in 1-1203 position (Table 3, Fig. 1). The most of the promoters (NM_001247420.1, XM_015224935.2, XM_015308168.1, XM_016725928.1,) of ZIP genes are positioned at 500 bp while some other are at 1200 bp (XM_019395528.1), 400 bp (XM_016650095.1, XM_009604423.3) and 200 bp (XM_028258694.1) position. However, no promoter was predicted in two ZIP protein (XM_016644574.1, XM_016650095.1) homologs (Table 3). The position of transcriptional start site (TSS) ranged from 201-1245, whereas the TATA-box was located from 884-1212 base positions (Table 3). We also found several AG and GT splice sites among the ZIP homologs across the plant species analyzed (Table 3).

**Table 3.** Organization of ZIP genes and position features.

| No. | Gene Accession | Exon number | Coding region | Promoter position | TATA-box position | Position of transcriptional start site (TSS) | Splice sites |
| --- | --- | --- | --- | --- | --- | --- | --- |
| 1 | NM_001247420.1 | 3 | 1-1053 | 500<br>(marginal prediction) | 911 | 946 | AG: 409, 444, 468, 658, 1012<br>GT: 694 |
| 2 | XM_015224935.2 | 3 | 1-1044 | 500<br>(marginal prediction) | 902 | 936 | AG: 397, 432, 659, 1003<br>GT: 685 |
| 3 | XM_015308168.1 | 3 | 1-1050 | 500<br>(marginal prediction) | 908 | 942 | AG: 397, 432, 655, 1009<br>GT: threshold |
| 4 | XM_009766342.1 | 3 | 37-1074 | No promoter predicted | 1212 | 1245 | AG: 229, 322, 679, 802, 1033, 1097, 1221<br>GT: 311, 547, 1359 |
| 5 | XM_016644574.1 | 3 | 163-975 | No promoter predicted | - | 261 | AG: 130, 223, 580, 703, 934<br>GT: 212, 448 |
| 6 | XM_016650095.1 | 3 | 122-984 | 400<br>(marginal prediction) | - | 252 | AG: 121, 130, 223, 375, 589, 712, 943<br>GT: 457 |
| 7 | XM_016725928.1 | 3 | 33-1024 | 500<br>(marginal prediction) | 884 | 918 | AG: 409<br>GT: Threshold |
| 8 | XM_009604423.3 | 3 | 163-984 | 400<br>(marginal prediction) | - | - | AG: 121, 223, 375, 589, 712<br>GT: 457 |
| 9 | XM_019395528.1 | 3 | 166-1203 | 1200<br>(highly likely prediction) | - | 201 | AG: 79, 349, 358, 451, 808, 931, 1162, 1216, 1332<br>GT: 440, 676, 1278 |
| 10 | XM_028258694.1 | 3 | 75-1088 | 200<br>(marginal prediction) | 1126 | 1159 | AG: 475, 784, 819, 983<br>GT: 514, 564, 732, 1234, 1365 |

**Fig. 1.**
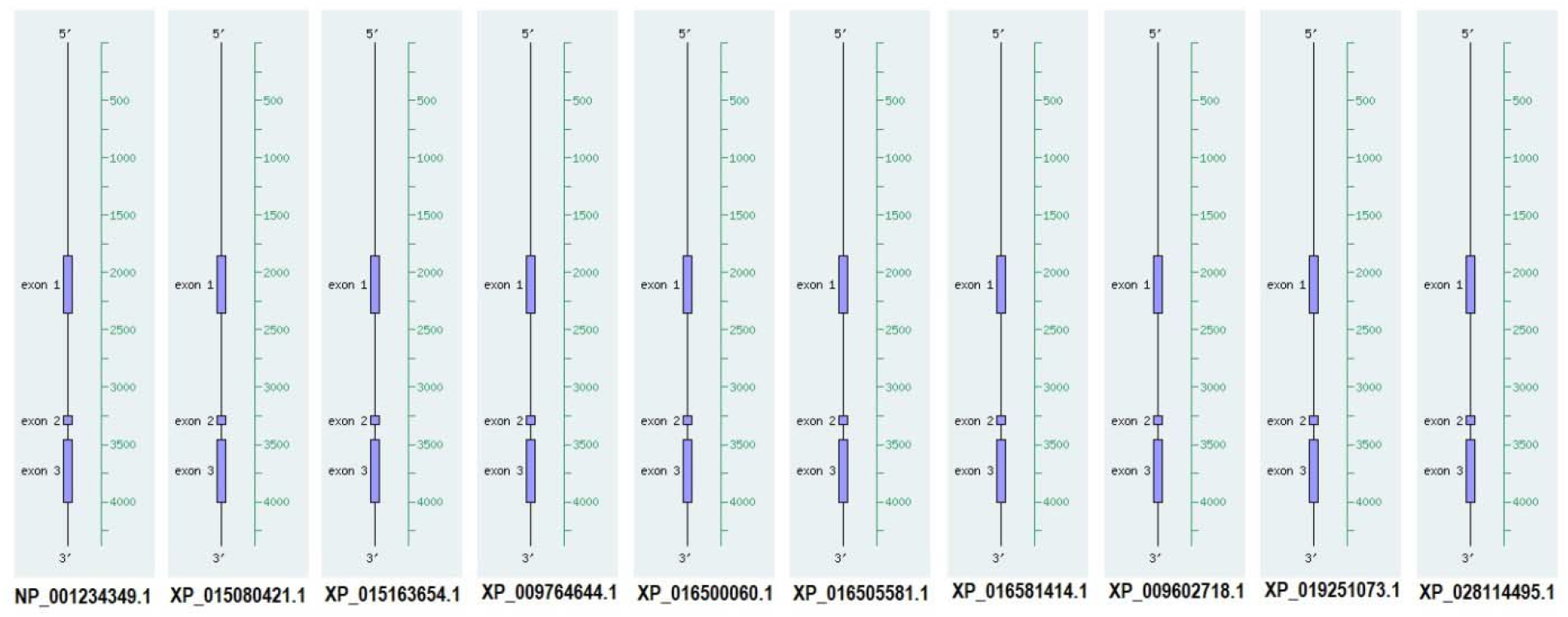
Gene organization of ZIP transporter homologs.

### Analysis of restriction enzyme and cutting position (Fig. 6)

We have searched for cutting positions and a list of restriction enzymes to explore the genetic information for future genome editing programs. All these ZIP transporters contain several cutting positions for several restriction enzymes (Fig. 2). Out of these restriction enzymes, *Bglll, Sacl* and *Nsil* are generally common across the ZIP transporter genes. The cutting position was ranged from 182-1442 base positions (Fig. 2).

**Fig. 2.**
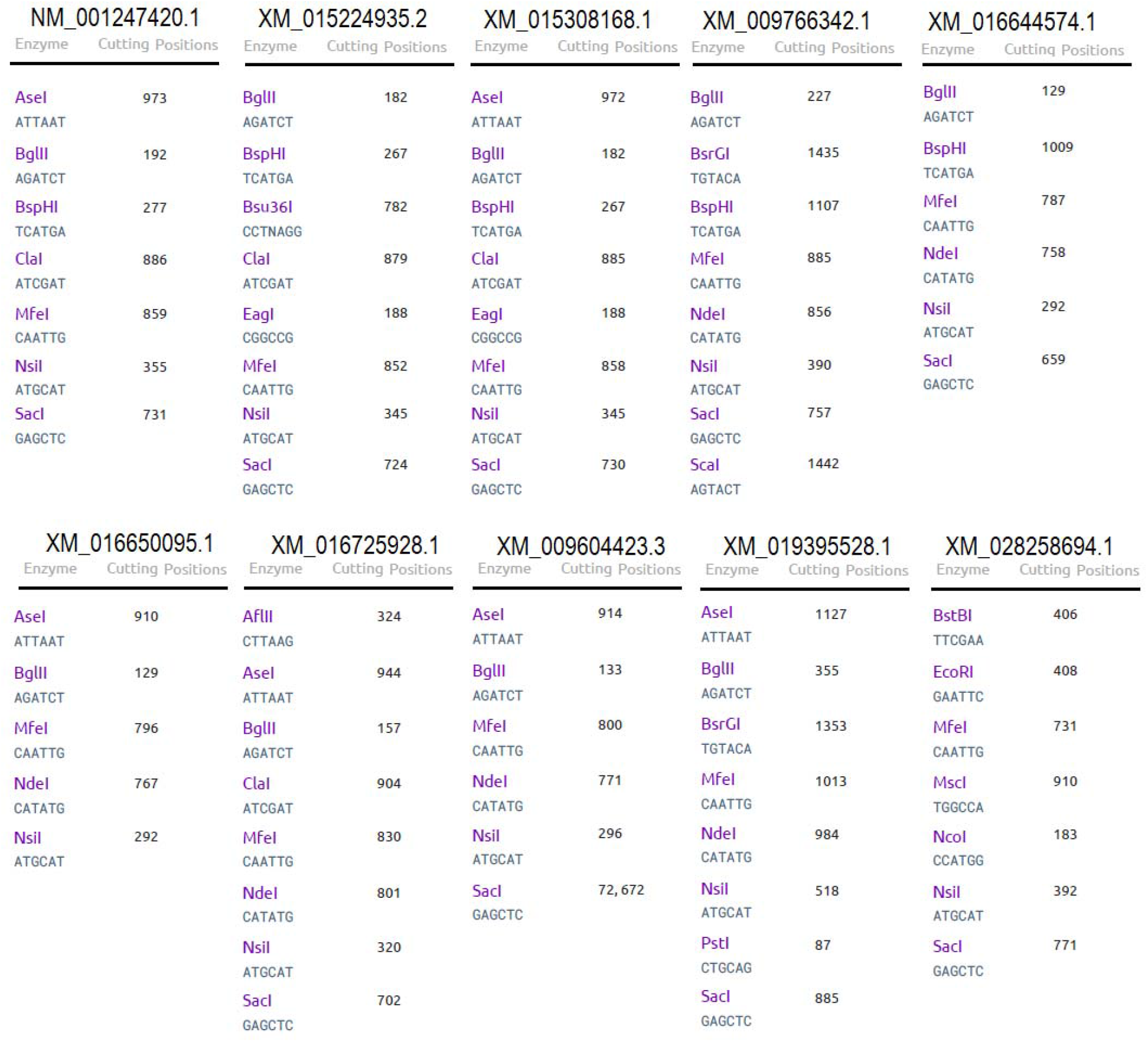
Restriction site and enzymes of ZIP transporter homologs.

**Fig. 3.**
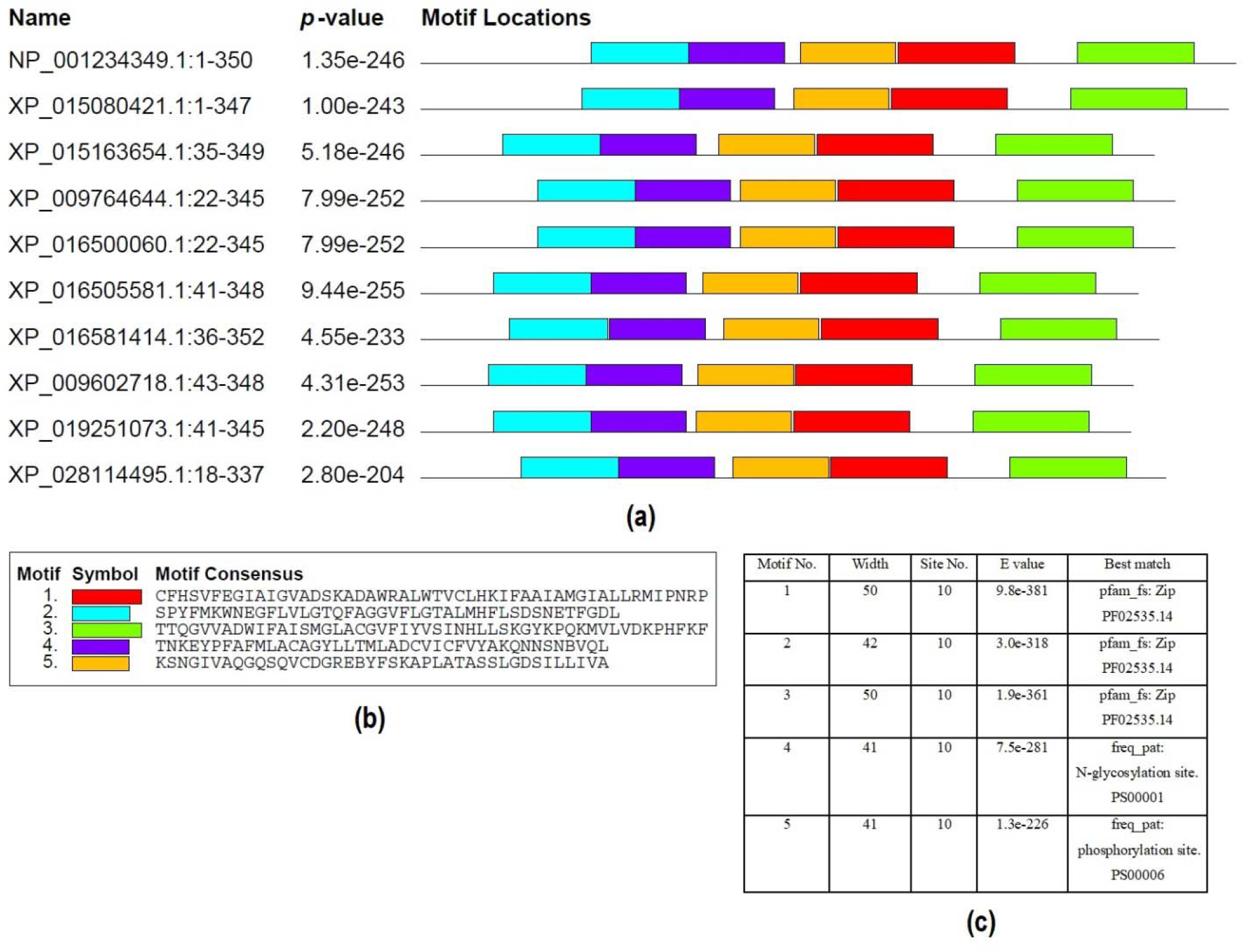
Most conserved five motifs of ZIP homologs detected by MEME tool.

**Fig. 4.**
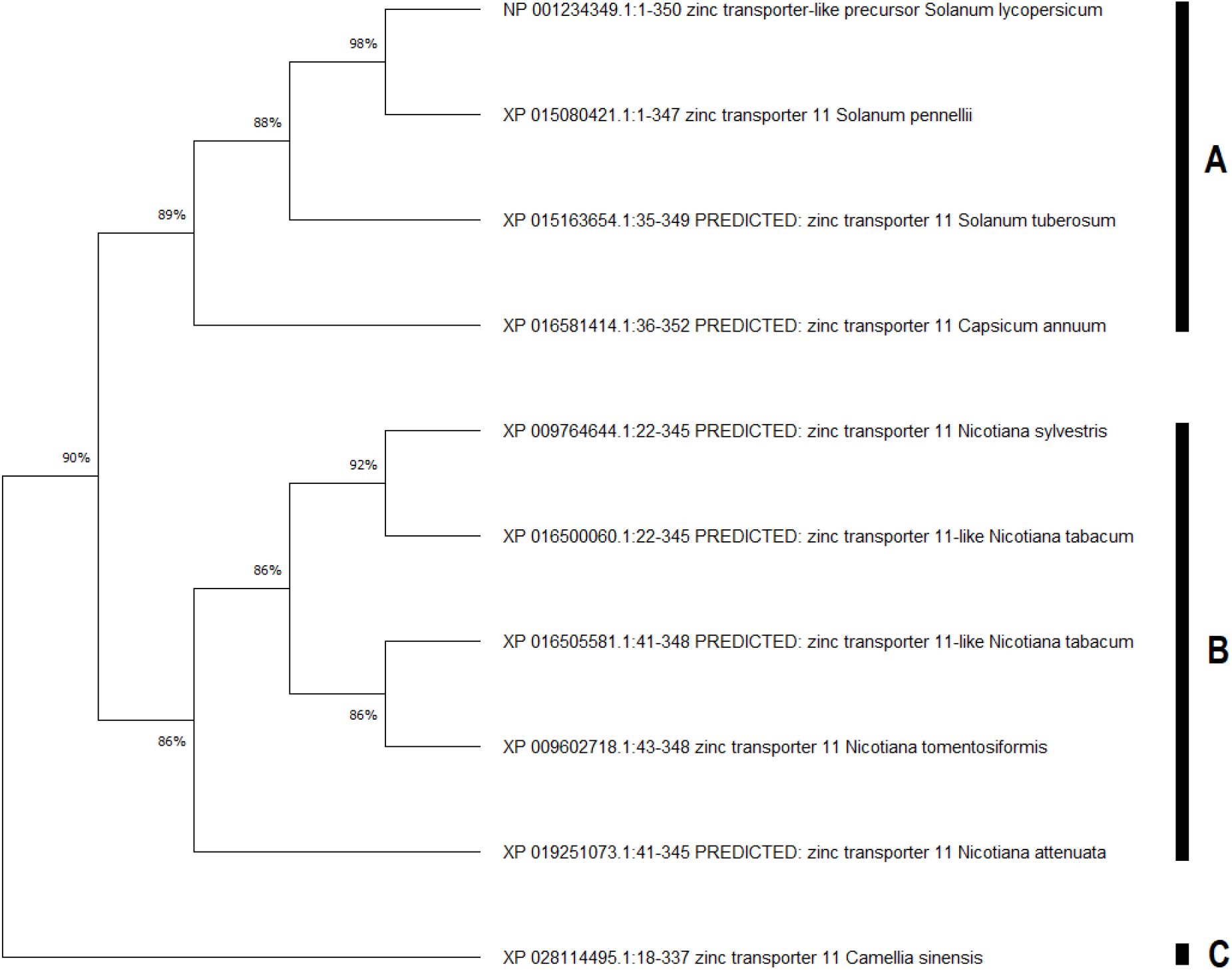
Phylogenetic tree of ZIP homologs using Mega 6. Statistical method: Maximum likelihood phylogeny test, test of phylogeny: bootstrap method, No. of bootstrap replications: 1000.

**Fig. 5.**
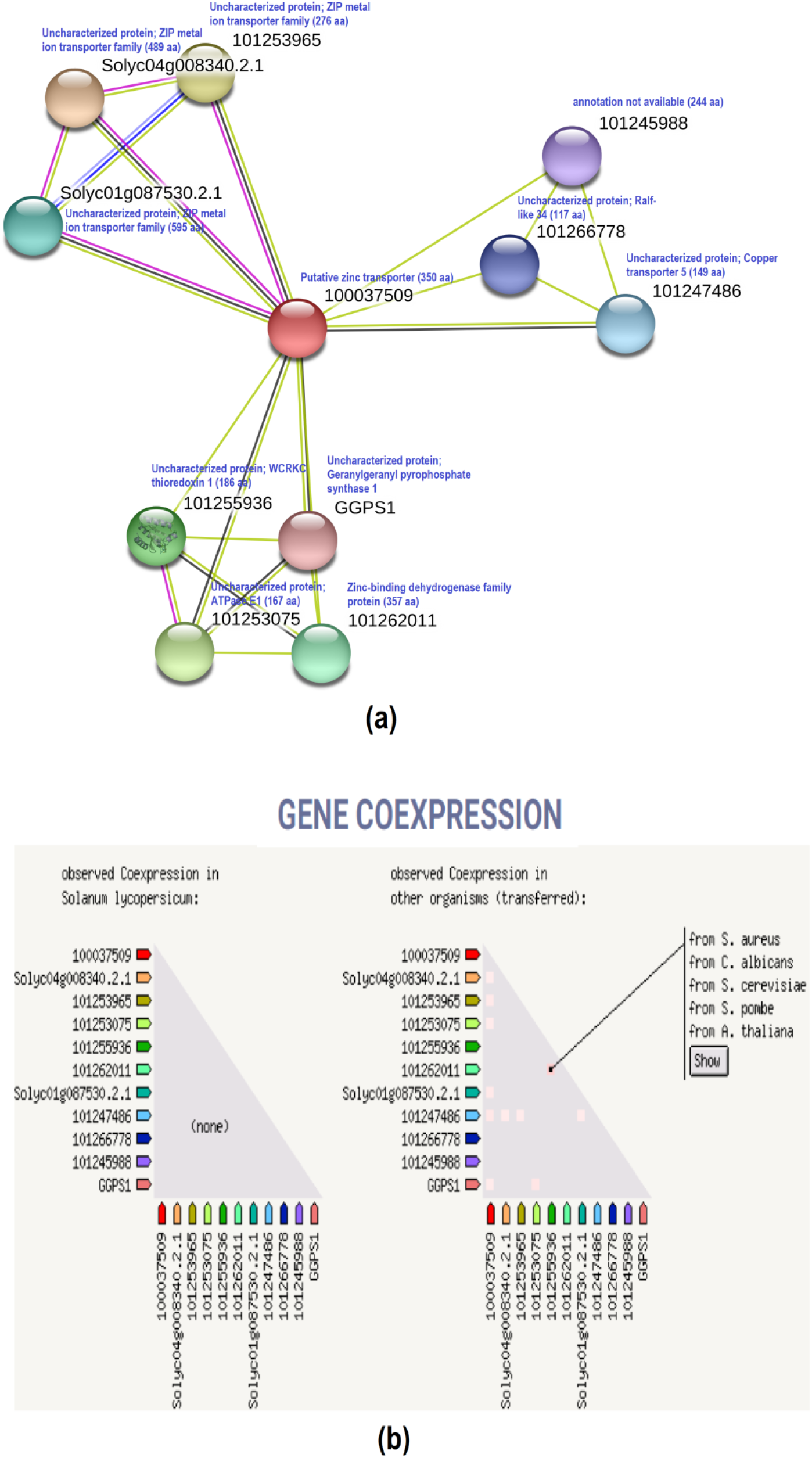
Predicted interaction partners of tomato ZIP transporter protein. Interactome was generated using Cytoscape for STRING data.

**Fig. 6.**
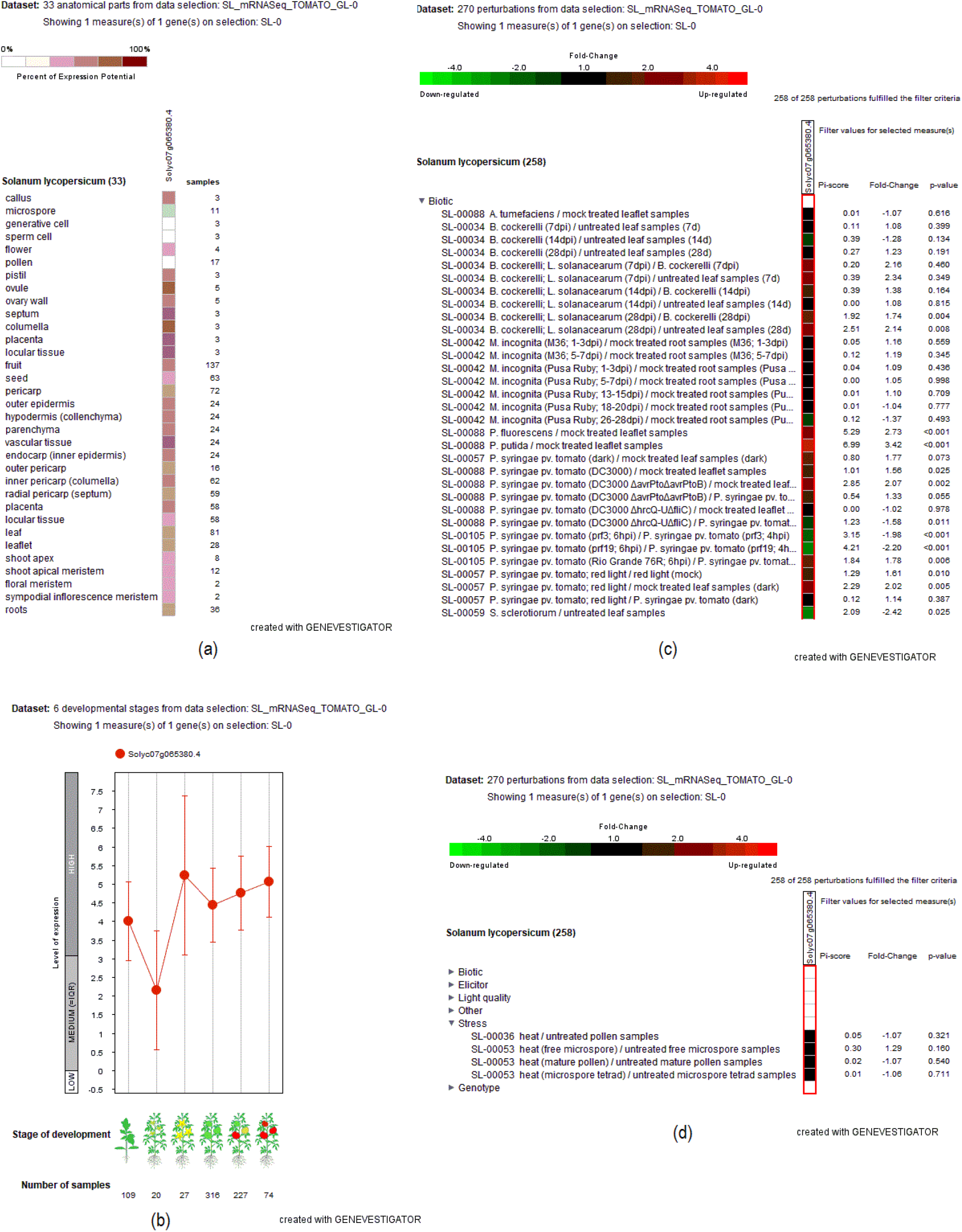
Co-expression of tomato ZIP transporter gene in different anatomical part, developmental stage and perturbations.

### Conserved motif, sequence similarities, and phylogenetic analysis

We have used the MEME tool to search for the five most conserved motifs in identified 10 ZIP protein homologs (Fig. 3). Motif 1, 2, 3, 4, and 5 showed 50, 42, 50, 41, and 41 long residues of amino acids, respectively. These five motifs were CFHSVFEGIAIGVADSKADAWRALWTVCLHKIFAAIAMGIALLRMIPNRP, SPYFMKWNEGFLVLGTQFAGGVFLGTALMHFLSDSNETFGDL, TTQGVVADWIFAISMGLACGVFIYVSINHLLSKGYKPQKMVLVDKPHFKF, TNKEYPFAFMLACAGYLLTMLADCVICFVYAKQNNSNBVQL, and KSNGIVAQGQSQVCDGREBYFSKAPLATASSLGDSILLIVA in the consecutive arrangement (Fig. 3). Out of these 5 motifs, motif 1-3 is best matched to PF02535.14 (ZIP), while motif 4 and 5 showed the best matching to PS00001 (N-glycosylation site) and PS0006 (phosphorylation site) sites (Fig. 3).

All also aligned all these 10 ZIP protein homologs to classify the regions of preserved (consensus) protein sequences. The ZIP protein homologs showed 94.6-100% sequence similarities among the species. The consensus sequence ranged from 70%-100% (Supplementary Fig. S1.). The phylogenetic was divided into two main groups based on tree topologies, such as A, B, and C (Fig. 4). In group A, ZIP transporter of *Solanum lycopersicum* showed 98%, 88% and 89% bootstrap with *Solanum pennellii, Solanum tuberosum* and *Capsicum annuum*, respectively (Fig. 4). Cluster B consisted of the 5 ZIP protein homologs, among which, *Nicotiana sylvestris* and *Nicotiana tabacum* showed highest 92% bootstrap value. The cluster C is solely placed with ZIP transporter of *Camelia sinensis* (Fig. 4).

### Predicted interaction partner analysis

Predicted interaction partner analysis was performed for tomato ZIP transporter (100037509). STRING showed ten interaction partners of 100037509 (350 aa), which include 101266778 (uncharacterized protein; Ralf-like 34), 101247486 (uncharacterized protein; copper transporter 5), 101245988 (annotation not available), Solyc01g087530.2.1 (uncharacterized protein; ZIP metal ion transporter family), Solyc04g008340.2.1 (uncharacterized protein; ZIP metal ion transporter family), 101253965 (uncharacterized protein; ZIP metal ion transporter family), 101255936 (uncharacterized protein; WCRKC thioredoxin 1, GGPSI (uncharacterized protein; geranylgeranyl pyrophosphate synthase 1), 101253075 (uncharacterized protein; ATPase EI), 101262011 (zinc-binding dehydrogenase family protein) proteins (Fig. 5a). Further, ZIP protein of *S. lycopersicum* showed co-expression with other protein members of the same species and other organisms (*S. aureus, C. albicans, S. cerevisiae, S. pombe, A. thaliana*) (Fig. 45).

The genvestigator analysis against SL_mRNASeq_Tomato_GL-0 showed co-expression data of tomato ZIP transporter (Solyc07g065380) in different anatomical parts, developmental stages and perturbations (Fig. 6a-c). In the anatomical part, the Solyc07g065380 showed the maximum potential of expression in the ovule, septum, columella, placenta, locular tissue, and vascular tissue (Fig. 6a). Also, Solyc07g065380 is highly predicted to be expressed during flower, fruit maturation, fruit ripening, fruit maturation, fruit initiation, and seedlings stages of development (Fig. 6b). Under perturbation analysis, we predicted the effect of biotic and abiotic stress interactions on Solyc07g065380 expression. It showed that Solyc07g065380 is highly upregulated (2-4 fold) in response to several pathogens, including *P. putida, B. cockerelli, P. fluorescens* and *P. syringae* (Fig. 6c). Although the data of abiotic stress is limited, Solyc07g065380 seems to be induced due to heat stress (Fig. 6d).

### Analysis of transmembrane helix in MTP1 proteins in different plant species

The outer membrane analysis of ZIP protein homologs exhibited variations in the localization of transmembrane strands and topology loops across the proteins as predicted by Philius transmembrane server (Fig. 7). Philius server displayed that only NP_001234349.1 and XP_015080421.1 proteins contain a signal peptide at the C-terminus end. In addition to these two ZIP proteins, XP_016581414.1 also displayed similar localization of 8 transmembrane helices in the protein sequences (Fig. 7). Further, XP_015163654.1, XP_009764644.1, XP_016500060.1 and XP_016505581.1 protein homologs without signal peptide showed identical localization of 7 transmembrane helices 3 (Fig. 7). The XP_009602718.1, XP_019251073.1 and XP_028114495.1 protein contain 7 transmembrane helices without any signal peptide (Fig. 7). Among the ZIP protein homologs, NP_001234349.1 and XP_015080421.1 showed the highest alpha helix, which is consistent with the presence of signal peptide (Supplementary Fig. S2). On average, the alpha helix of all these ZIP proteins ranged from 24.06% to 31.12% of the protein structure. In addition, the extended strand contains 21.60% to 28.12% of the structural organization of ZIP proteins (Supplementary Fig. S2). As expected, random coil contains stands roughly around 50% locations of all ZIP proteins. None of these ZIP protein homologs displayed the presence of 3_10_ helix, Pi helix, beta bridge, beta turn, bend region and ambiguous state in structure (Supplementary Fig. S2).

**Fig. 7.**
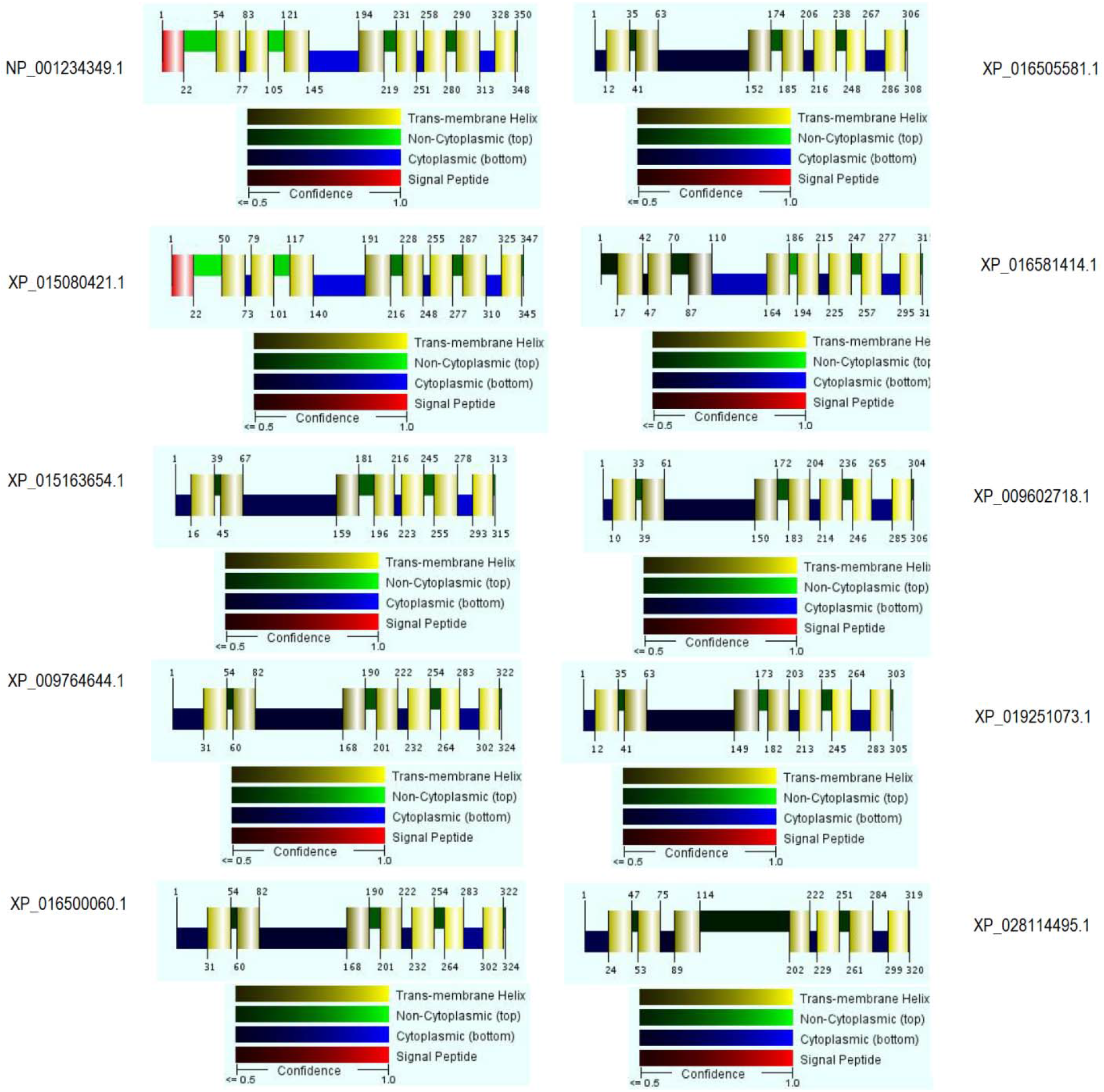
Prediction of transmembrane helix of ZIP protein homologs.

## Discussion

Characterization of a gene is of great interest to further accelerate the wet-lab experiments on plant stress. This *in silico* work led to the identification of 10 ZIP protein homologs among nine plant species. MSA shows these proteins are 98.5-100% coverage of similarity with 94.6-100% matching of consensus sequences among the different ZIP proteins. Most ZIP proteins are predicted to have eight potential transmembrane domains and a similar membrane topology in which the amino- and carboxy-terminal ends of the protein are located on the outside surface of the plasma membrane (Guerinot 2000). Our *in silico* analysis showed the existence of 7-8 transmembrane helices in all 10 ZIP protein homologs connected to ZIP Zinc transporter (PF02535). All of these ZIP protein homologs are localized in chromosome 7 and the plasma membrane.

The position and organization of the coding sequence of a gene is considered to be a critical factor in predicting evolutionary relations and functional genomics potentialities. In this study, all ZIP gene homologs among the nine plant species showed 3 exons, suggesting that these ZIP genes are phylogenetically closer to each other. Localization of promoter plays an essential part in CRISPR-Cas9 and other genome editing studies in plant science. Our analysis suggests that the promoter of *ZIP* gene homologs are highly positioned at 400-500 bp. Additionally, we explored the position of TSS and splice sites of ZIP homologs, which are crucial in understanding the transcriptional and post-transcriptional modification of mRNA. Restriction site and specificity of restriction enzymes are essential before transformation studies on a particular gene of interest. In this study, several cutting sites and respective restriction enzymes have been revealed, which will be useful to develop transgenic tomato or other plants associated with Zn homeostasis. In general, it is known that genes without intron have recently evolved (Deshmukh et al. 2015), which is not the case for ZIP gene homologs, as evident in our analysis. ZIP transporters are not involved in cellular Zn homeostasis but also regulate the adaptive growth response in plants (Coninx et al. 2019).

Conserved motifs are identical sequences across species that are maintained by natural selection. A highly conserved sequence is of functional roles in plants and can be a useful start point to start research on a particular topic of interest (Wong et al. 2015). Among the 10 ZIP protein homologs, we searched for five motifs using the MEME tool. Out of the five motifs, three motifs mainly matched with the ZIP zinc transporter family. We also observed the N-glycosylation site in motif 4, which is in agreement with the previous report on the conserved features of NRAMP transporter (Curie et al. 2000). The relation of motif 5 with the phosphorylation site may be attributed to the release of Zn from intracellular stores leading to phosphorylation kinases and activation of signaling pathways as previously reported (Thingholm et al. 2020). The presence of common and long conserved residues pinpoints that ZIP homologs may possess highly conserved structures between species. Additionally, this information can be targeted for sequence-specific binding sites and transcription factor analysis.

In phylogenetic analysis, we clustered the tree in 3 sub-groups. According to the tree, ZIP protein of *S. lycopersicum* showed the closest relationship (98% bootstraps) with *S. pennellii*, while *S. tuberosum* and *C. annuum* also existed with the cluster of A. Consistently, ZIP protein homologs of five tobacco species clustered within B, suggesting the close evolutionary emergence from a common ancestor. It also appears that the ZIP transporter of *S. lycopersicum* is relatively distantly related to *C. sinensis* over the evolutional trends. However, it is not yet reported whether this Solyc07g065380 putative Zn transporter is also involved in Zn uptake in other closely related plant species. Thus, our results might infer a functional relationship ZIP sequences in Zn or other metal uptake across different plant species.

Interactome map and co-expresion analysis were performed using the Solyc07g065380 in the STRING platform. In the interactome map, tomato putative zinc transporter (100037509) seemed to be associated with several interaction partners linked to uncharacterized ZIP metal ion transporter family. As a result, these findings might be useful to characterize several ZIP metal transporters in plants. In plants, ZIP family members have also been characterized in plants involved in metal uptake and transport, including Zn (Kavitha et al., 2015). However, studies also reported that Zn homeostasis is closed associated with P-type ATPase heavy metal transporters (HMA). Both HMA2 and HMA4 were reported to be involved with Zn homeostasis in Arabidopsis (Hussain et al., 2004). Our further search on the coexpression of Solyc07g065380 suggests that this particular ZIP transporter is highly possible to coexpressed with several genes of *S. lycopersicum* as well as *A. thalina*. Overall, this interactome findings might provide essential background for functional genomics studies of Zn uptake and transport in plants.

The potentiality of expression of a gene in different conditions is a crucial factor in the genome editing program. The *in silico* analysis of expression in the Genevestigator platform showed interesting outputs concerning the expression of Solyc07g065380 in different anatomical, perturbations, and developmental stages. Although root is the primary focus of ZIP transporter, this Genevestigator platform suggested the maximum potential of expression in ovule, septum, columella, placenta, locular tissue, and vascular tissue. In wet lab experiences, several CDF and ATPase family transporters were shown root-specific expression regulating Zn and Cu homeostasis in plants (Desbrosses-Fonrouge et al. 2005). Further, Solyc07g065380 is predominantly expressed in several development stages during maturation, although the expression potential during seedlings development is crucial for Zn uptake and Zn-deficiency tolerance in tomato. Besides, the expression of Solyc07g065380 seemed to be upregulated in response to several pathogens and heat stress. Although the data of stress is limited in the Genevestigator platform, it implies that Solyc07g065380 is highly active in response to perturbation. Given this potentiality of Solyc07g065380, this study further advances our knowledge to elucidate the uptake and mobilization of Zn and other metals in plants.

In this study, all of the identified sequences of ZIP proteins demonstrated acidic character having the pI value of around 6 along with the stable instability index and positive hydropathicity. The protein length of 10 ZIP proteins was 305-350. Several studies reported the length of amino acid residues in the ZIP family transporter from 309–476 (Guerinot, 2000). Computational prediction of the secondary structure further reveals the presence of the signal peptide in ZIP protein homologs of *S. lycopersicum* and *S. pennellii*. Signal peptides function to prompt a cell to translocate the protein, usually to the cellular membrane. Thus, it suggests that the ZIP protein of these two Solanum species carries enormous potentiality in the Zn uptake system in plants. The highest % of the alpha helix in these two ZIP protein homologs is in accordance with the higher transmembrane domains as integral membrane proteins are predominantly located in alpha-helices. These findings may provide insights into the protein architecture and particular function.

## Conclusion

In conclusion, these computational studies analyzed 10 ZIP protein homologs in different plant species. The analysis showed similar physicochemical properties, gene organization, and conserved motifs related to the Zn ion transport. Sequence homology and phylogenetic tree of ZIP proteins showed the closest evolutionary relationship of *S. lycopersicum* with *S. pennellii*. In addition, the interactome map displayed the co-expression of Solyc07g065380 with a number of closely related genes involved with the ZIP metal ion transporter family. In addition, Solyc07g065380 does have a high potentiality to express in several organs at different developmental stages subjected to biotic and heat stress in the Genevestigator platform. This *in silico* analysis reveals several gene features, such as promoter position, TSS, restriction sites, and enzymes, which will be useful for functional genomics and transgenic studies related to Zn uptake in plants. The unique feature of having signal peptide in Solyc07g065380 and other structural organization of protein further confirm the potentiality of this putative Zn transporter. These findings will provide basic theoretical knowledge for future studies to better understand the function and features of gene/protein related to Zn homeostasis in various plants.

## Acknowledgment

I would like to acknowledge the accompaniment of my little daughters, Arya and Alina without whom it was hard to get rid of the depression and to concentrate on this work in COVID-19 pandemic.

---

**Supplementary Fig. S1.**
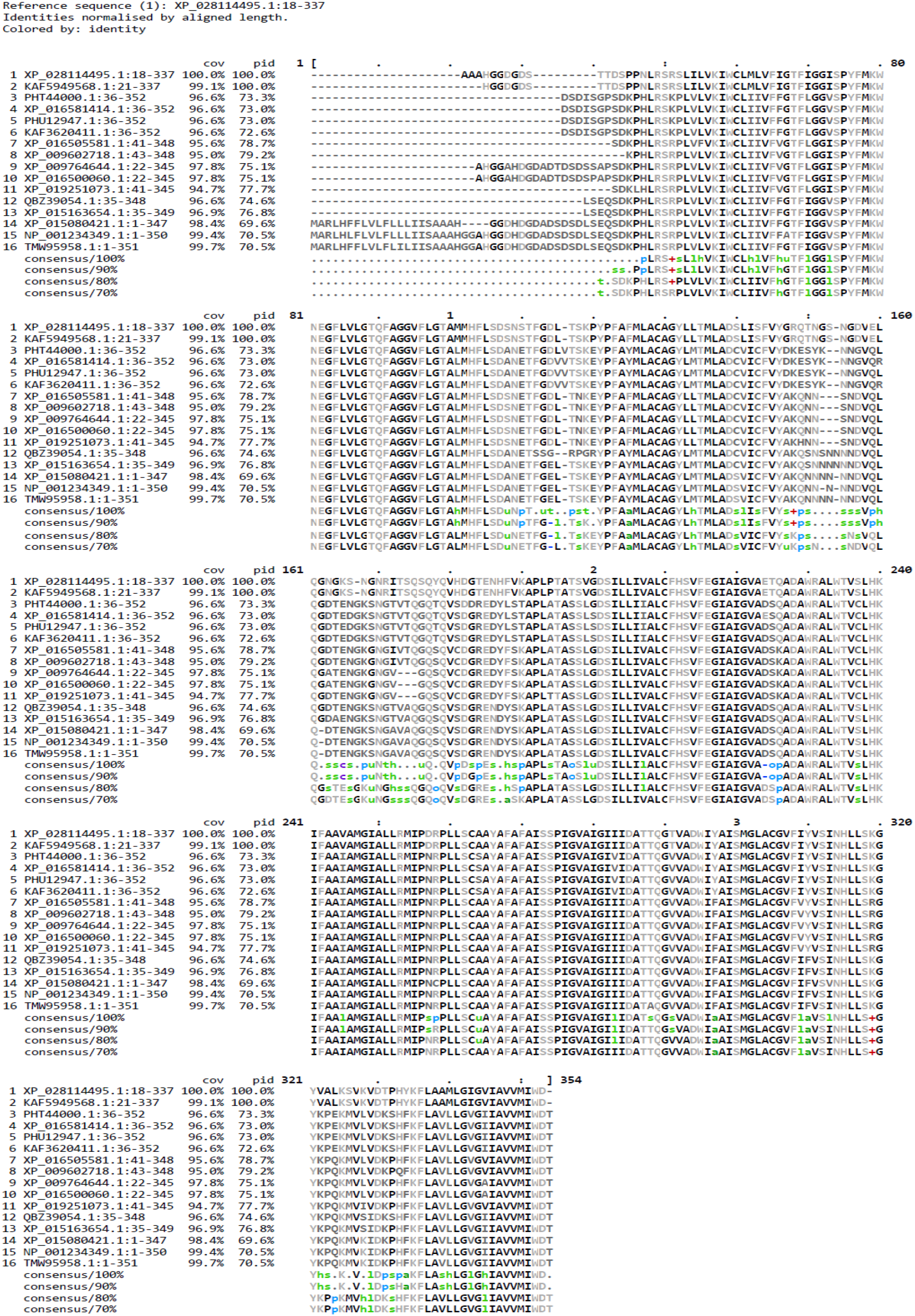
Multiple sequence alignment of ZIP protein homologs.

**Supplementary Fig. S2.**
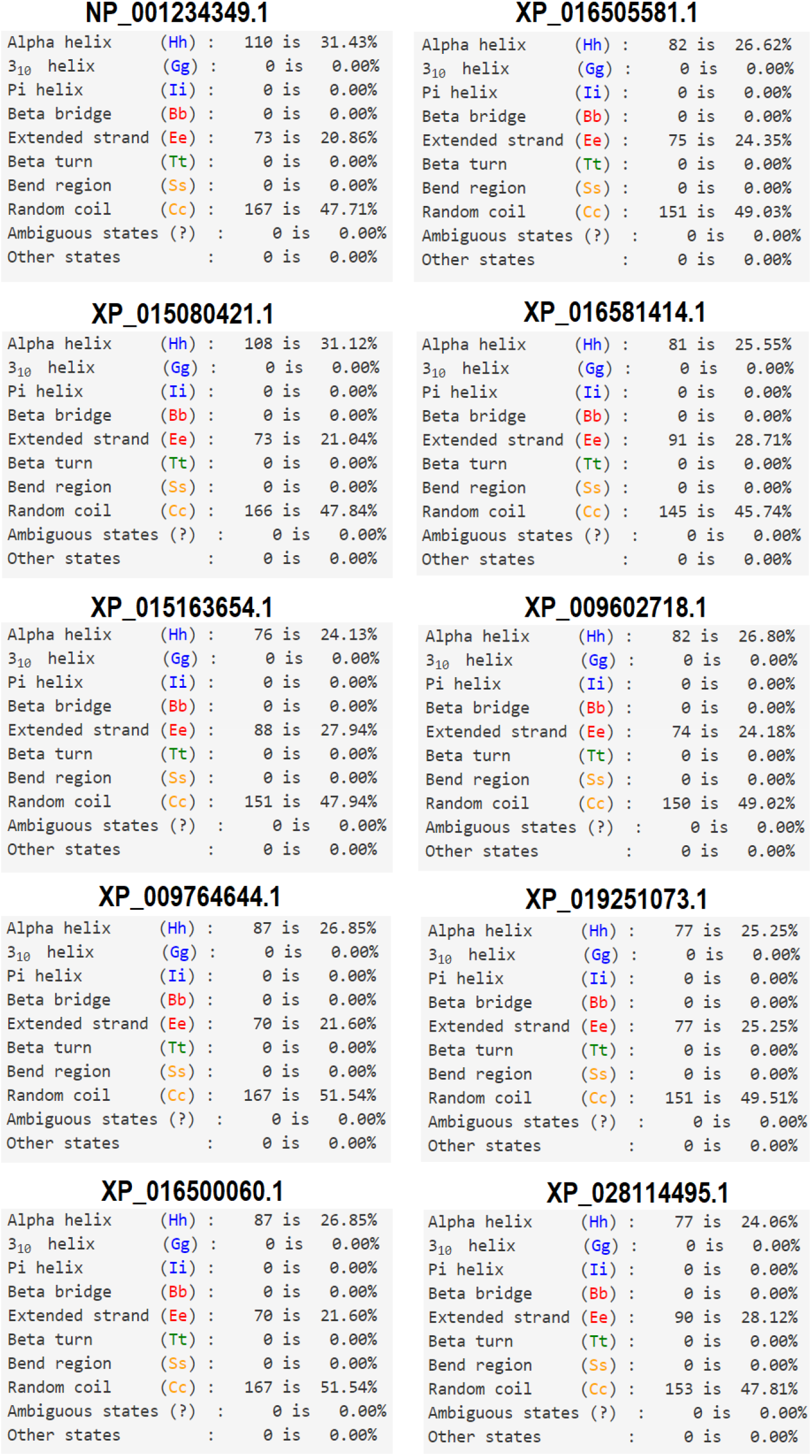
Description of secondary structure of ZIP protein homologs.

